# A genome-wide association study to identify the genetic *loci* underlying carbapenem resistance in *Acinetobacter baumannii*

**DOI:** 10.64898/2025.12.16.694545

**Authors:** Gurprit Sekhon, Balvinder Singh

## Abstract

Carbapenem-resistant *Acinetobacter baumannii* (CRAB) is a major public health threat and a key contributor to the global antimicrobial resistance (AMR) crisis. This study applied a bacterial genome-wide association study (bGWAS) framework to identify genetic factors beyond the canonical carbapenemase OXA-23, that may contribute to carbapenem resistance. *A. baumannii* isolates with publicly available whole-genome sequencing (WGS) data and carbapenem-specific antimicrobial susceptibility testing (AST) results were compiled (n = 1,601). Unitigs were extracted from *de novo* assemblies, and single-nucleotide polymorphisms (SNPs) were called from raw sequence reads mapped to seven selected reference genomes. The population was highly clonal and comprised ten lineages, including three predominantly resistant ST2 lineages. The known CRAB-associated *loci*, including OXA-23, the RND efflux pump regulator AdeL, and an OXA-66 substitution (Leu167Val), were readily identified. Novel resistance-associated mutations were detected in outer-membrane proteins, including porins (OprD, DcaP) and proteins (PqiB, LptD) involved in lipopolysaccharide (LPS) biogenesis. Notably, the detection of OprD and DcaP was reference-dependent, indicating that a multi-reference bGWAS framework improves identification of resistance-associated variants that may be missed in single-reference analyses. Structural mapping of the identified mutations suggested plausible effects on protein function.

## INTRODUCTION

Antimicrobial resistance (AMR) is a global public health crisis [1]. *Acinetobacter baumannii* has emerged as a major nosocomial bacterial pathogen that contributes substantially to the global AMR burden [2, 3]. Carbapenem-resistant *A. baumannii* (CRAB) typically exhibits a multidrug-resistant (MDR) phenotype, severely limiting therapeutic options [4]. CRAB increasingly compromises the effectiveness of carbapenem antibiotics, which are among the last-line options for treating MDR infections in clinical settings [5]. The World Health Organization (WHO) has designated CRAB as a critical-priority pathogen due to its high disease burden and potent ability to disseminate resistance genes [6].

Carbapenem resistance is predominantly mediated by carbapenem-hydrolyzing class B (metallo-*β*-lactamases) and class D (oxacillinases) β-lactamases [7, 8]. In *A. baumannii*, OXA-51–like β-lactamases are intrinsic and ubiquitous but exhibit only weak carbapenemase activity. In contrast, acquired OXA-23–family enzymes are the most widespread clinically relevant class-D carbapenemases [9]. However, even potent carbapenemases such as *OXA-23* do not fully account for the carbapenem resistance observed in many clinical isolates [10]. This has motivated investigations into additional carbapenem resistance mechanisms. In particular, reduced outer-membrane permeability due to porin disruption and increased efflux activity *via* resistance–nodulation–division (RND) pumps have been implicated in modulating carbapenem susceptibility [11, 12].

Genome-wide association studies (GWAS) have provided important insights into antimicrobial resistance determinants across diverse bacterial species [13, 14]. A hypothesis-free bacterial GWAS framework is well suited to identify genetic variants contributing to carbapenem resistance in *A. baumannii*, including those beyond established mechanisms. To enable such an analysis, publicly available *A. baumannii* genomes with matched carbapenem-specific antimicrobial-susceptibility testing (AST) data were curated, and unitig- and SNP-based genotype–phenotype association testing was conducted. To the best of our knowledge, this represents the first CRAB-focused GWAS with a large sample size (n = 1,601).

## RESULTS

### The genome dataset and quality control

A schematic overview of the study workflow is shown in Supplementary Figure S1. Initially, 1814 publicly available *A. baumannii* genomes with carbapenem-specific AST data were identified. Of these, only 1565 isolates had raw sequence reads and were assembled *de novo*, while the genome assemblies for the remaining 249 isolates were retrieved from NCBI. The quality of raw sequence reads for all but eight isolates, which contained non-biological reads (adapter contamination), is summarized in Supplementary Figure S2. Quality control was performed on the resulting genome assemblies (n=1806) in two stages (Supplementary Figure S3). In stage 1, 153 assemblies were excluded for poor reference coverage (<80%) or low Contig-N50 (<20 kb). In stage 2, an additional 52 assemblies were excluded for low genome completeness (<99%) or high contamination (>5%), yielding a final set of 1601 assemblies for downstream analysis (Supplementary Figure S3). For these genome assemblies (1601), metadata such as geographic origin, isolation year, isolation source, and SRA/Assembly accessions are provided in Supplementary Information (“*Metadata.tsv*”).

### Population structure and phylogenetic analysis

Bayesian clustering resolved the population into ten lineages (BAPS_1–BAPS_10), corresponding to clades I–X in Figure 1. BAPS_5 (n = 747), BAPS_4 (n = 305), BAPS_8 (n = 149), BAPS_2 (n = 120), and BAPS_9 (n = 116) contained more than 100 isolates. BAPS_2, BAPS_5, and BAPS_9 were predominantly resistant, whereas BAPS_4 and BAPS_8 were mainly susceptible (Figure 1). The first ten MDS axes corresponded to the BAPS lineages (Supplementary Figure S4), and explained more than 80% variation in population structure (Supplementary Figure S5a). First two MDS axes separated the major resistant and susceptible lineages (Supplementary Figure S5b).

**Figure 1.**
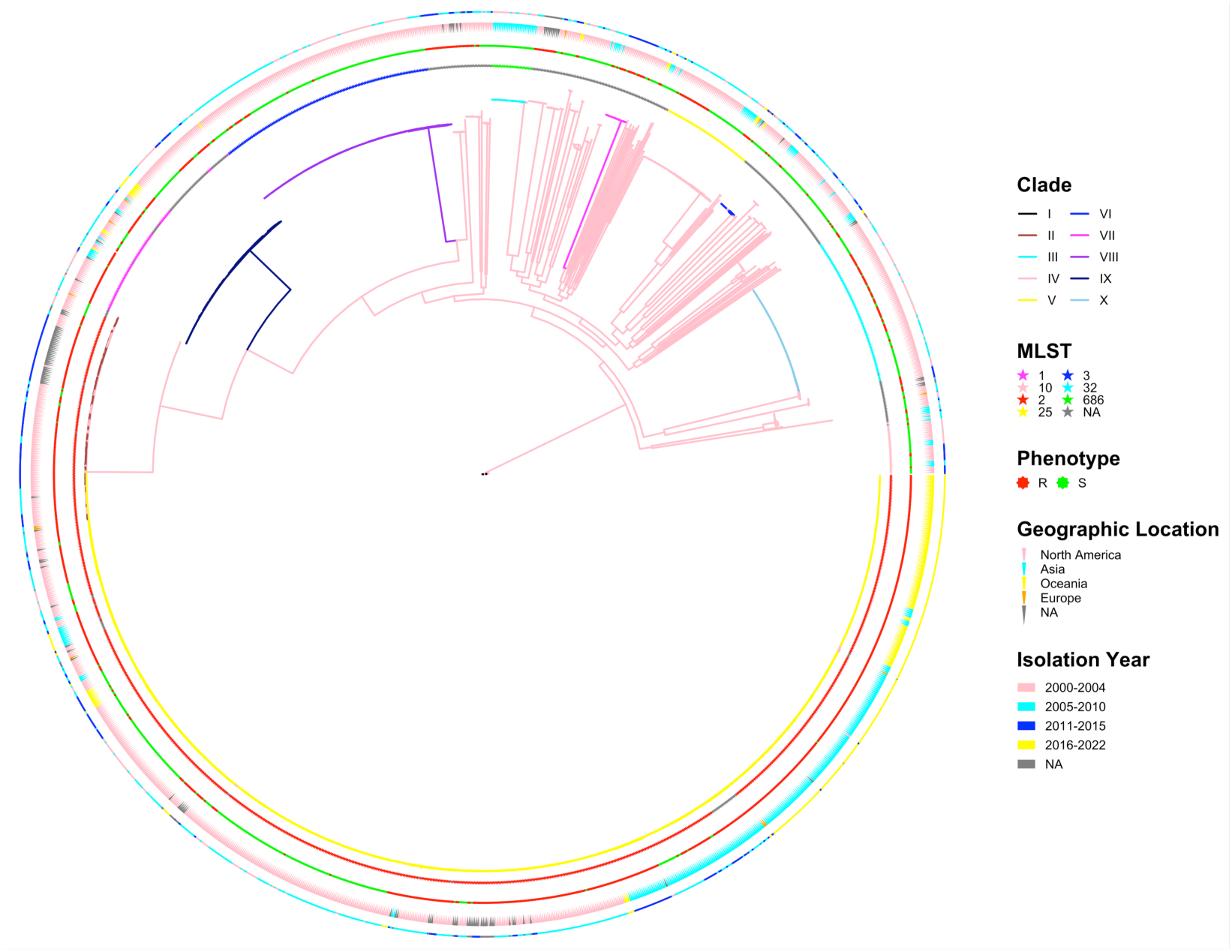
Population structure and phylogenetic analysis. Recombination-corrected maximum-likelihood phylogeny based on the core genome SNPs in 1,601 *A. baumannii* isolates. The phylogeny is colored according to BAPS-defined lineages. The innermost ring denotes sequence type (ST), followed by rings representing carbapenem phenotype, geographic origin and isolation year, respectively. (BAPS, Bayesian analysis of population structure)

The resistant lineages BAPS_2 and BAPS_5 formed a single clade in the phylogenetic tree (Figure 1). Consistently, the first MDS axis accounted for both BAPS_2 and BAPS_5 (Supplementary Figure S4) and these lineages clustered together in the MDS1-MDS2 plot (Supplementary Figures S5b). By comparison, the resistant lineage BAPS_9 formed a distinct clade on the phylogenetic tree (Figure 1) and also clustered separately from BAPS_2 and BAPS_5 in MDS analysis (Supplementary Figure S5b). Sequence type (ST) assignments were consistent with the inferred population structure. Isolates from the resistant lineages (BAPS_2, BAPS_5 and BAPS_9) belonged to ST2, the most prevalent CRAB sequence type, whereas the susceptible lineages BAPS_8 and BAPS_4 corresponded to ST3 and ST25, respectively (Figure 1).

The separation between resistant and susceptible lineages was apparent in the phylogeny (Figure 1). BAPS_2, BAPS_9, and BAPS_8 were composed almost exclusively of isolates from the USA. Within the largest resistant lineage, BAPS_5, isolates from Asia and Oceania clustered separately from those originating in the USA (Figure 1). BAPS_8 contained a substantial number of isolates from earlier isolation years (2000–2004). In addition, a smaller resistant lineage, BAPS_10 (n = 88, ST3), comprised predominantly resistant isolates from the USA, sampled between 2002 and 2010 (Figure 1).

### Genome-wide association study

The results from unitig- and SNP-based GWAS across seven reference genomes are shown in Figure 2. Total number of unitigs and mean number of SNPs identified across reference genomes were 1,971,690 and 257,803±8,353, respectively. The effect of population structure correction is illustrated by the observed versus expected *P* value distributions for the unitig (Figure 2a) and SNP (Figure 2b) analyses. In both QQ plots, the observed *–log_10_(P)* followed the expected null distribution across the lower range (approximately *–log₁₀(P)* < 2), indicating adequate control of inflation (Figures 2a and 2b). Using Bonferroni-adjusted significance thresholds of 5 × 10⁻⁸ for unitigs and 2.5 × 10⁻⁷ for SNPs, 999 significant unitigs and an average of 606 ±64 significant SNPs across reference genomes were identified. The association results for unitigs and SNPs are presented in Figures 2c–2p, wherein each set of two panels show results for a single reference genome.

**Figure 2.**
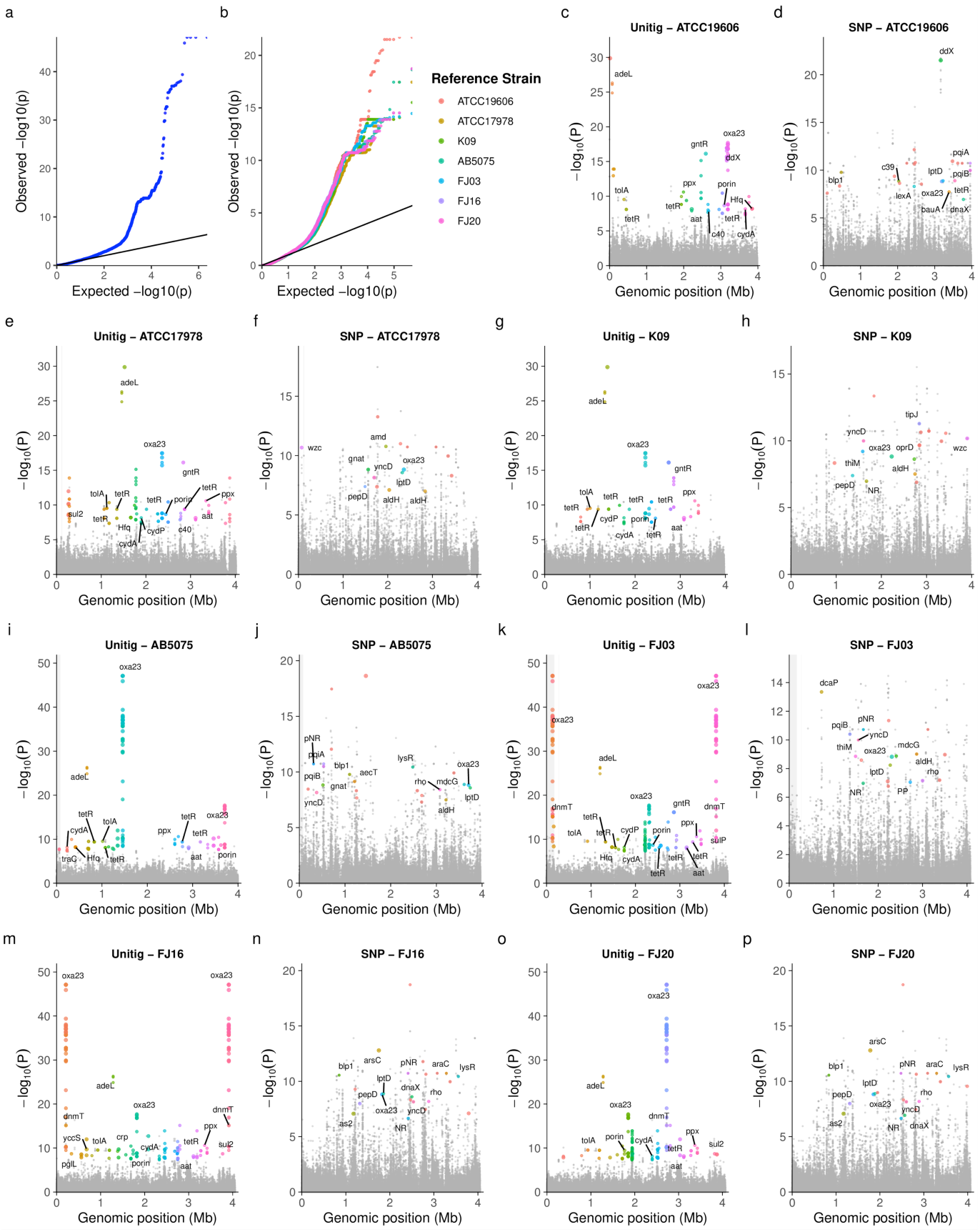
Significant genetic *loci* identified by genome-wide association analyses. (a,b) QQ plots comparing observed and expected *P* values for unitig- and SNP-based GWAS, respectively. (c–p) Manhattan plots showing genome-wide association signals from unitig and SNP-based analyses for each of the seven reference genomes.

Both the acquired carbapenemase, *OXA-23* and the intrinsic OXA-51 family member, *OXA-66* were annotated with identical labels across reference genomes (Figure 2c-2p). The genes were reassigned to their families based on sequence alignment. *OXA-23* showed markedly stronger association signals than *OXA-66* (Figures 2i, 2k, 2m, 2o). ATCC19606, ATCC17978, and K09 encoded only *OXA-66*, whereas the remaining reference genomes contained both *OXA-23* and *OXA-66* (Figures 2). A known OXA-66 mutation showed significant association across reference genomes (Figure 2d, 2f, 2h, 2j, 2l, 2n, 2p). The Leu167Val substitution results from C499G nucleotide change and has been reported to increase carbapenemase activity [15, 16]. Leu167 forms part of the enzyme binding pocket, and substitution to Valine has been suggested to facilitate carbapenem binding [16, 17].

The LysR-type transcription factor *adeL* showed significant association across reference genomes [Figure 2c, 2h, 2i, 2k, 2m, 2o]. AdeL is the repressor of RND efflux pump adeFGH in *A. baumannii* and its overexpression leads to almost twofold increase in imipenem MIC [12]. Therefore, RND efflux pumps regulators, including AdeL, are attractive drug targets to combat AMR in *A. baumannii* [18].

Leu382Pro substitution in outer-membrane (OM) porin OprD was found to be significant (Figure 2h). OprD belongs to OM carboxylate channel (OCC) family implicated in carbapenems permeation across outer membrane in Gram-negative bacteria. Four subfamilies (OCCAB1-4) have been characterized in *A. baumannii*, with OCCAB1 having the widest pore size [19]. The identified OprD shares 35% and 25% sequence identity with OCCAB4 (PDB ID: 5dl8) and OCCAB2/3 (PDB ID: 5dl6, 5dl7), respectively and each of these shows more than 90% query coverage. No significant sequence similarity was observed with OCCAB1. The C-terminal region containing Leu382 showed no detectable sequence homology (Figure 3a). The AlphaFold-predicted structure showed an RMSD of 1.03 Å to the crystal structure (PDB ID: 5dl8), and the substituted residue lies within the extracellular vestibule loop (Figure 3b).

**Figure 3.**
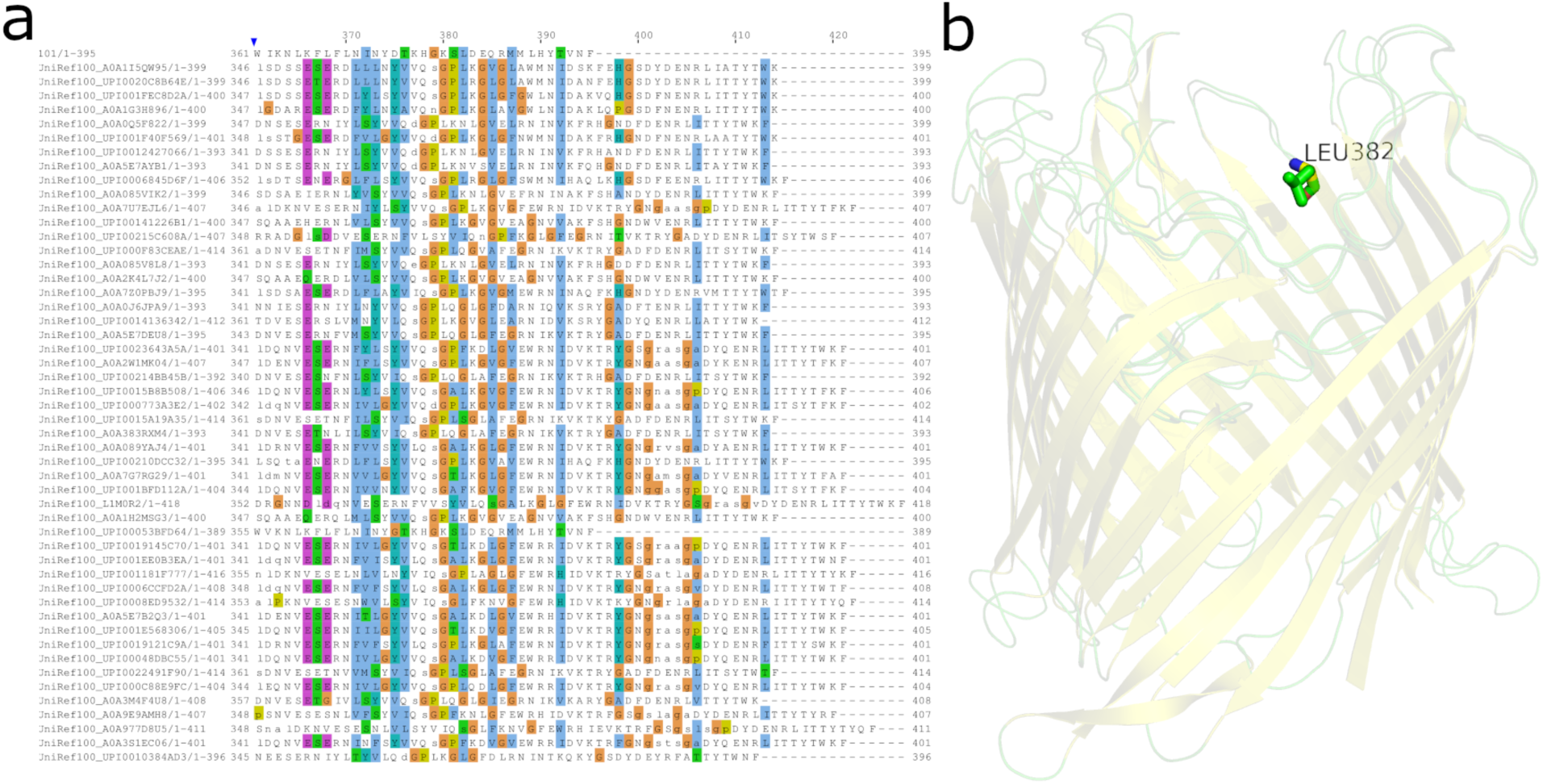
The Leu382Pro mutation identified in the outer-membrane porin OprD. (a) Excerpt of AlphaFold-generated multiple-sequence alignment (MSA) showing the C-terminal region of OprD having the affected residue. (b) In the AlphaFold-predicted structure, Leu382 lies within the extracellular vestibule of the OprD porin, suggesting a potential impact on substrate passage.

A single residue deletion in DcaP (ΔSer338) was significantly associated with carbapenem resistance (Figure 2l). DcaP is a highly expressed outer-membrane porin during *A. baumannii* infection in rodent models and has been proposed to mediate the cellular uptake of clinically relevant small *β*-lactamase inhibitors [20]. Ser338 showed limited sequence conservation among homolog proteins (Figure 4a).

**Figure 4.**
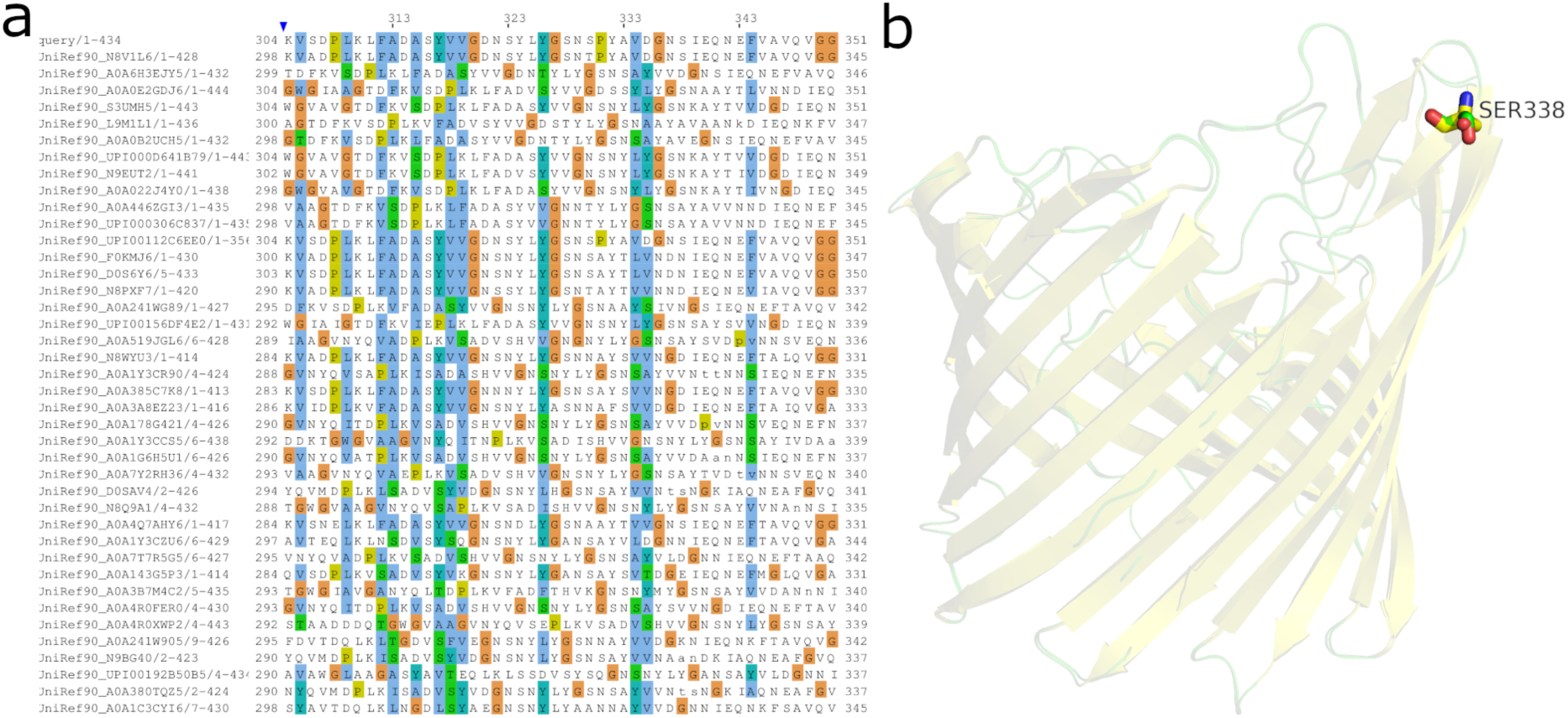
The ΔSer338 mutation identified in the outer-membrane porin, DcaP. (a) Excerpt of AlphaFold-generated MSA shows limited conservation at residue 338. Ser338 (shown as sticks) is part of extracellular loop region in the AlphaFold-predicted structure.

The identified DcaP sequence was found to be 97% identical to the corresponding sequence in crystal structure (PDBID: 6EUS), with more than 80% query coverage. Structural superposition of the AlphaFold-predicted model and the crystal structure yielded an RMSD of 0.2 Å. The Ser338 is localized to an extracellular loop region (Figure 4b) and it may potentially affect the substrate permeation.

Multiple substitutions were detected in outer-membrane proteins involved in lipopolysaccharide (LPS) biogenesis. These included mutations in PqiB, which is involved in LPS transport (Figures 2d, 2j, 2l), and LptD, an essential component of the LPS assembly machinery (Figures 2d, 2f, 2j, 2l, 2n, 2p) [21]. Structural superposition of the AlphaFold-predicted LptD model onto the homologous crystal structure (PDB ID: 8H1S; RMSD 1.8 Å; 30% Sequence Identity) showed that the substituted residues cluster within a *β*–turn region of the integral membrane barrel domain (Figure 5a). In addition, significant associations were observed in transcriptional regulators such as members of LysR, TetR, and GntR families (Figure 2). Notably, a TetR-family regulator carried a significant mutation (Arg24Cys, Figure 2d). Genomic context analysis showed that this regulator is located upstream of a TonB-dependent receptor. Structural analysis of the Alphafold-predicted TetR model indicated that the substituted Arg24 residue is located within the DNA-binding domain [Figure 5b].

**Figure 5.**
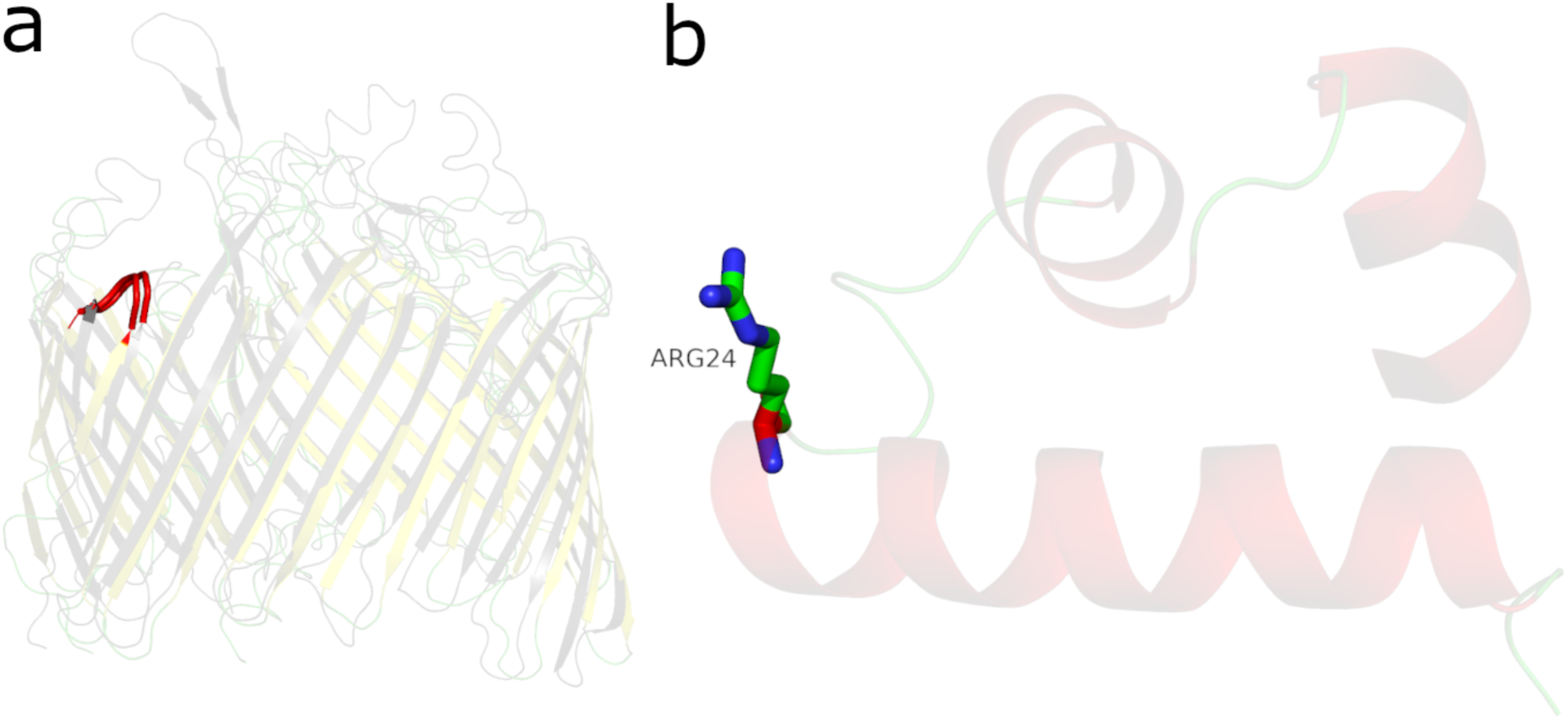
Mutations identified in LptD and TetR-family transcriptional regulator. (a) Structural superposition of the AlphaFold-predicted LptD model (Yellow) onto the crystal structure (Grey, PDB ID: 8H1S) shows substituted residues clustered in a *β*-turn region (Red). (b) AlphaFold-predicted structure of the TetR-family regulator with the Arg24Cys substitution (shown in sticks) mapped to the DNA-binding domain.

The unitig- and SNP-based GWAS consistently identified OXA-23, OXA-66 (Leu167Val), and AdeL as the strongest resistance-associated signals across the population (Figures 2c–2p). Significant associations were also detected for outer-membrane proteins LptD and PqiB across multiple reference genomes (Figures 2d, 2f, 2h, 2j, 2l, 2n, 2p). Beyond these broadly conserved associations, several *loci* exhibited reference-dependent detection patterns. Specifically, OprD and DcaP achieved genome-wide significance when analyzed with respect to reference genomes K09 (Figure 2h) and FJ03 (Figure 2l), respectively. A complete list of mutations and association statistics is provided in Supplementary Information (Table S1).

## DISCUSSION

Carbapenem-resistant *A. baumannii* (CRAB) remains a major clinical threat, and resistance is commonly attributed to the acquired carbapenemase, *OXA-23*. However, the resistance observed in many CRAB isolates cannot be fully explained by carbapenemase activity alone [10]. An elucidation of additional resistance determinants may help more accurately define the genetic basis of carbapenem resistance. Genome-wide association studies (GWAS) provide a robust, hypothesis-free approach for identifying such genotype–phenotype correlations. To date, comprehensive carbapenem-focused GWAS analyses in *A. baumannii* remain limited. In this study, a large-scale bacterial GWAS integrating genomic and carbapenem susceptibility data was conducted on 1,601 publically available *A. baumannii* isolates. A key strength of the present analytical framework was the combined use of a multi-reference SNP-based GWAS and a reference-independent unitig approach. This multi-reference strategy enabled the identification of resistance-associated variants whose detectability and statistical significance depended on the choice of reference genome(s).

Clonal population structure is a major confounding factor in bacterial GWAS [22]. Accordingly, the population structure of the GWAS dataset was characterized prior to association testing. Bayesian clustering resolved ten distinct lineages (BAPS_1–BAPS_10), with BAPS_5 and BAPS_8 representing predominantly resistant and susceptible lineages, respectively, and BAPS_2 & BAPS_9 constituting smaller resistant clusters (Figure 1). Multidimensional scaling (MDS) supported this lineage structure (Supplementary Figure S4) and the first ten MDS axes captured more than 80% of the population structure variance (Supplementary Figure S5a). These findings are consistent with the strong clonality reported in other bacterial pathogens [23]. Major lineages were clearly separated along the first two MDS axes, with BAPS_2 and BAPS_5 clustering together, BAPS_9 positioned more distantly, and BAPS_8 forming a distinct susceptible lineage [Supplementary Figure S5b].

These relationships were further corroborated by the recombination-filtered phylogeny (Figure 1). Concordantly, BAPS_5, BAPS_2, and BAPS_9 were classified as ST2, whereas BAPS_8 corresponded to ST3, and isolation year metadata showed that the earliest isolates were predominantly assigned to BAPS_8 (Figure 1). These results confirm strong lineage structure within the dataset and underscore the importance of accounting for clonal population structure in bacterial GWAS.

The present study identified OXA-23 as the strongest resistance-associated *locus*, consistent with previous reports implicating this carbapenemase as a primary driver of carbapenem resistance in *A. baumannii* [10]. In addition, the transcriptional repressor AdeL and an OXA-66 variant (Leu167Val) were significantly associated with carbapenem resistance, in agreement with earlier studies linking both AdeL-mediated efflux regulation [12] and OXA-66 variants [16] to altered carbapenem susceptibility.

Novel resistance-associated mutations were identified in the outer-membrane proteins. These include membrane porins OprD (Leu382Pro) & DcaP (ΔSer338), and proteins involved in lipopolysaccharide (LPS) transport (PqiB) and assembly (LptD). OprD is known to mediate carbapenem uptake [24], whereas DcaP has been implicated in the uptake of *β*-lactamase inhibitors [20]. Preliminary sequence and structural analysis indicated that the substituted residues in OprD and DcaP are moderately conserved; however, their localization within the extracellular vestibules of the respective channels suggests that these mutations could alter substrate permeability (Figures 3 and 4).

Novel resistance-associated mutations were identified in PqiB (Gly432Asp, Lys474Asn, Asn430Ser, Ala440Pro, Leu407Phe) and LptD (Asn280dup, Val290Ile, Asp281Gly, Ser279_Asn280insGlyPheSer, Gln421His, Ile284Val, Ile284Ser). *PqiB* is a member of the MCE superfamily, found exclusively in Gram-negative bacteria, and participates in lipopolysaccharide (LPS) transport across the periplasm [25]. LptD is an essential *β*-barrel component of the LPS assembly machinery at the outer membrane [26]. In *A. baumannii*, disruption of LptD is tolerated, although deletion mutants exhibit impaired growth and increased membrane permeability [27]. Notably, most of the identified LptD mutations cluster within a *β*-turn region positioned toward the membrane surface (Figure 5a). The variations observed in PqiB and LptD may perturb LPS biogenesis, thus affecting membrane permeability. Together these porin- and LPS-associated mutations highlight the role of outer-membrane permeability in shaping antimicrobial susceptibility in Gram-negative bacteria [28].

Transcriptional regulators such as LysR*-*, TetR-, and GntR-family proteins were significantly associated with resistance (Figure 2), indicating the potential contribution of transcriptional regulation to the resistance phenotype. Among these, a novel mutation in the TetR-family regulator (Arg24Cys) was found to be located within the DNA-binding domain (Figure 5b), suggesting a potential effect on regulation of downstream TonB*-*dependent outer-membrane receptor proteins. This observation is consistent with the well-established role of TetR regulators in mediating antibiotic resistance [29]. In addition, several resistance-associated mutations were detected in hypothetical or poorly characterized proteins (Supplementary Table 1), highlighting gaps in current understanding of CRAB biology and prioritizing candidate *loci* for follow-up studies.

The present GWAS identified novel resistance-associated variants in outer-membrane porins (OprD, DcaP), LPS-associated proteins (PqiB, LptD), transcriptional regulators, and less well-characterized proteins, thereby expanding the genetic landscape linked to carbapenem resistance in *A. baumannii*. This study is constrained by reliance on heterogeneous publicly available AST data and would have benefited from quantitative MIC measurements rather than binary resistance phenotypes. Consequently, the identified associations are based on genomic signals and require experimental validation.

## METHODS

### Acquisition, preprocessing, and quality control

Using the BV-BRC command-line interface (*v1.035*) [30], publicly available *A. baumannii* genomes were queried, yielding 11,824 genome identifiers. These identifiers were subsequently queried for experimental AST results, producing 19,413 resistant and 5,344 susceptible phenotype entries. AST records were filtered for carbapenems (doripenem, ertapenem, imipenem, and meropenem), generating a carbapenems-specific dataset of 862 resistant and 841 susceptible genomes. AST results for a subset of genomes (n = 45) indicated resistance to some carbapenems but susceptibility to others; therefore, these genomes were excluded from analysis. This resulted in a preliminary cohort of 817 resistant and 796 susceptible isolates. An additional 201 carbapenem-resistant isolates from a recent molecular epidemiological study from the Oceanic region were incorporated [31], yielding a final dataset of 1,814 genomes (1,018 resistant, 796 susceptible). SRA/Assembly accessions, geographic origin, isolation year, and isolation source metadata were retrieved for the 1814 isolates.

Sequence reads were retrieved from the Sequence Read Archive (SRA) using Sra-tools (*v3.1.0*) [32]. Raw reads were quality-filtered and adapter-trimmed using Fastp (*v0.22.0*) [33], and quality summaries were assessed using MultiQC (*v1.14*) [34]. Trimmed reads were assembled *de novo* using SPAdes (*v3.15.5*) [35]. For isolates lacking raw SRA data, genome assemblies were downloaded directly from NCBI [*36*]. Assembly quality was evaluated in two stages using QUAST (*v5.2.0*) and CheckM (*v1.2.2*) [37, 38]. QUAST was used to assess assembly statistics relative to the AB5075 reference genome, and assemblies with >80% reference coverage and N50 >20,000 bp were retained. CheckM was then applied to estimate completeness and contamination based on lineage-specific marker genes; assemblies with <99% completeness or >5% contamination were excluded. Only genomes passing both quality filters were included in downstream analyses.

### Phylogenetic Analysis

Core genome SNP alignments were generated using Snippy (*v4.6.0*) with AB5075 as the reference genome. Variant sites were extracted using SNP-sites (*v2.5.1*) [39]. Recombinant regions were detected and masked using Gubbins (v3.3.1) [40]. The recombination-corrected alignment was used to infer a maximum-likelihood phylogeny in RAxML (*v8.2.12*) under the *GTR+Γ* substitution model with 500 bootstrap replicates [41]. The resulting tree was visualized and annotated in R (*v4.3.3*) using the ggtree (*v3.10.1*) and ggtreeExtra (*v1.12.0*) packages [42, 43]. The population structure was inferred using RhierBAPS (*v1.1.4*) [44, 45]. Multilocus sequence typing (MLST) was performed using mlst (*v2.23.0*).

### Multi-reference Strategy

To mitigate reference bias and capture pan-genome variation, raw sequence reads were independently aligned to seven representative *A. baumannii* genomes spanning clinical, environmental, and epidemiologically diverse backgrounds. The selected references included ATCC19606 (susceptible clinical isolate; BioSample SAMN12389466), ATCC17978 (clinical meningitis isolate; BioSample SAMN02604331), K09 (environmental isolate from soil; BioSample SAMN12769618), AB5075 (hypervirulent clinical isolate; BioSample SAMN31681936) [46], and three isolates from a recent CRAB epidemiological study in the Oceania region: FJ03, FJ16, and FJ20 (BioSamples SAMN38749311, SAMN37733145, SAMN37736705, respectively) [31].

### Genome-wide Association Analysis

Raw sequence reads and genome assemblies (SRA data not available) were processed with Snippy (*v4.6.0*) to generate reference-aligned BAM files for each of the seven reference genomes. Variant calling was performed using joint genotyping in GATK (*v4.5.0.0*) [47]. Missing genotypes were imputed using the major allele at the corresponding site. Bifrost, wrapped as unitig-caller (*v1.3.0*), was used to extract unitigs [48]. GWAS was performed using the factored spectrally transformed linear mixed model (FaST-LMM) implemented in Pyseer (*v1.3.11*) [49]. The model incorporated geographic origin and isolation year as fixed covariates, and a phylogeny-derived kinship matrix as a random effect. SNPs were annotated with SnpEff (v5.0e) [50]. Bonferroni correction was applied using the number of unique unitig and SNP patterns evaluated in the respective analyses.

## DECLARATIONS

### 1. Ethics approval

Not Applicable.

### 2. Consent for publication

Not Applicable.

### 3. Data

All data used in present study is publically available. The SRA/Assembly accessions (“Metadata.tsv”), BASH and R scripts used for workflow automation and data analysis are available at https://github.com/sekhongurprit/GWAS.

### 6. Authors’ contributions

BS and GS designed the study. GS carried out the work, analyzed the data, and prepared the draft. BS and GS reviewed the subsequent drafts and made necessary changes before finalizing the manuscript.

## Supporting information

Metadata (Supplementary File)

## 7. Acknowledgements

GS gratefully acknowledges the Indian Council of Medical Research (ICMR) for funding the research fellowship during this project, and the Council of Scientific & Industrial Research–Institute of Microbial Technology (CSIR-IMTECH) for providing the laboratory space and computational resources necessary to carry out this work.

## 8. Impact

This genome-wide association study of carbapenem-resistant *Acinetobacter baumannii* identifies candidate resistance *loci* in outer-membrane porins (OprD, DcaP) and proteins (LptD, PqiB) involved in lipopolysaccharide (LPS) biogenesis, highlighting the contribution of Gram-negative outer-membrane permeability to carbapenem resistance. By employing a multi-reference mapping strategy, the analysis mitigates reference-dependent detection biases and enables identification of resistance-associated variants that may be missed in single-reference approaches. Preliminary sequence and structural analyses prioritize these *loci* for further mechanistic investigation. Importantly, this study demonstrates that well-controlled bacterial GWAS can detect both canonical and previously unreported resistance-associated signals, even in highly clonal bacterial pathogens.

## Supplementary Information

**Figure S1.**
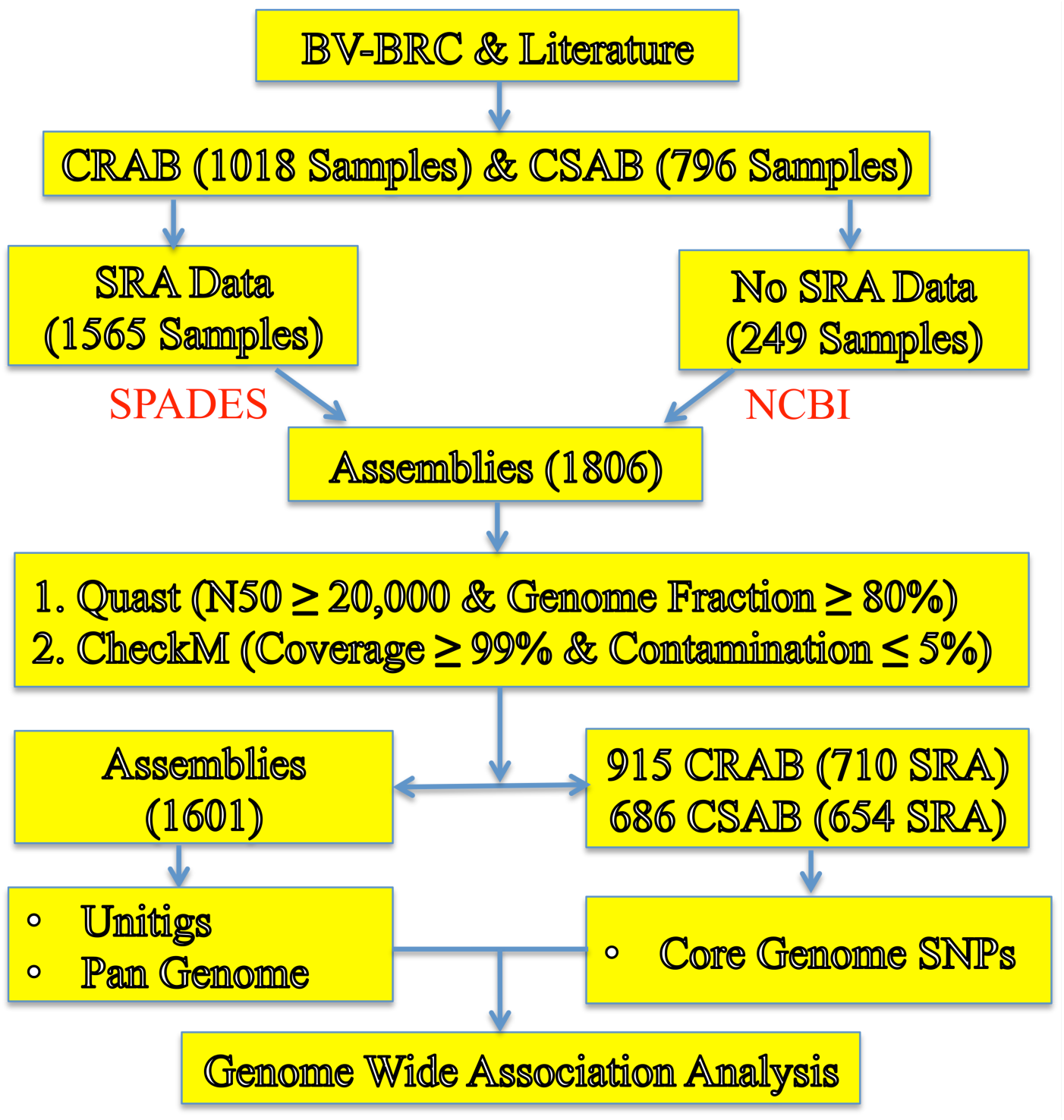
Schematic overview of the GWAS pipeline. The workflow includes acquisition of sequencing data, genome assembly, quality control, unitig extraction, SNP calling, and genotype–phenotype association analysis. (BV-BRC, Bacterial and Viral Bioinformatics Resource Center; CRAB, Carbapenem Resistant *A. baumannii*; CSAB, Carbapenem Susceptible *A. baumannii*; SPADES, *de novo* assembly by SPAdes; NCBI, National Center for Biotechnology Information)

**Figure S2.**
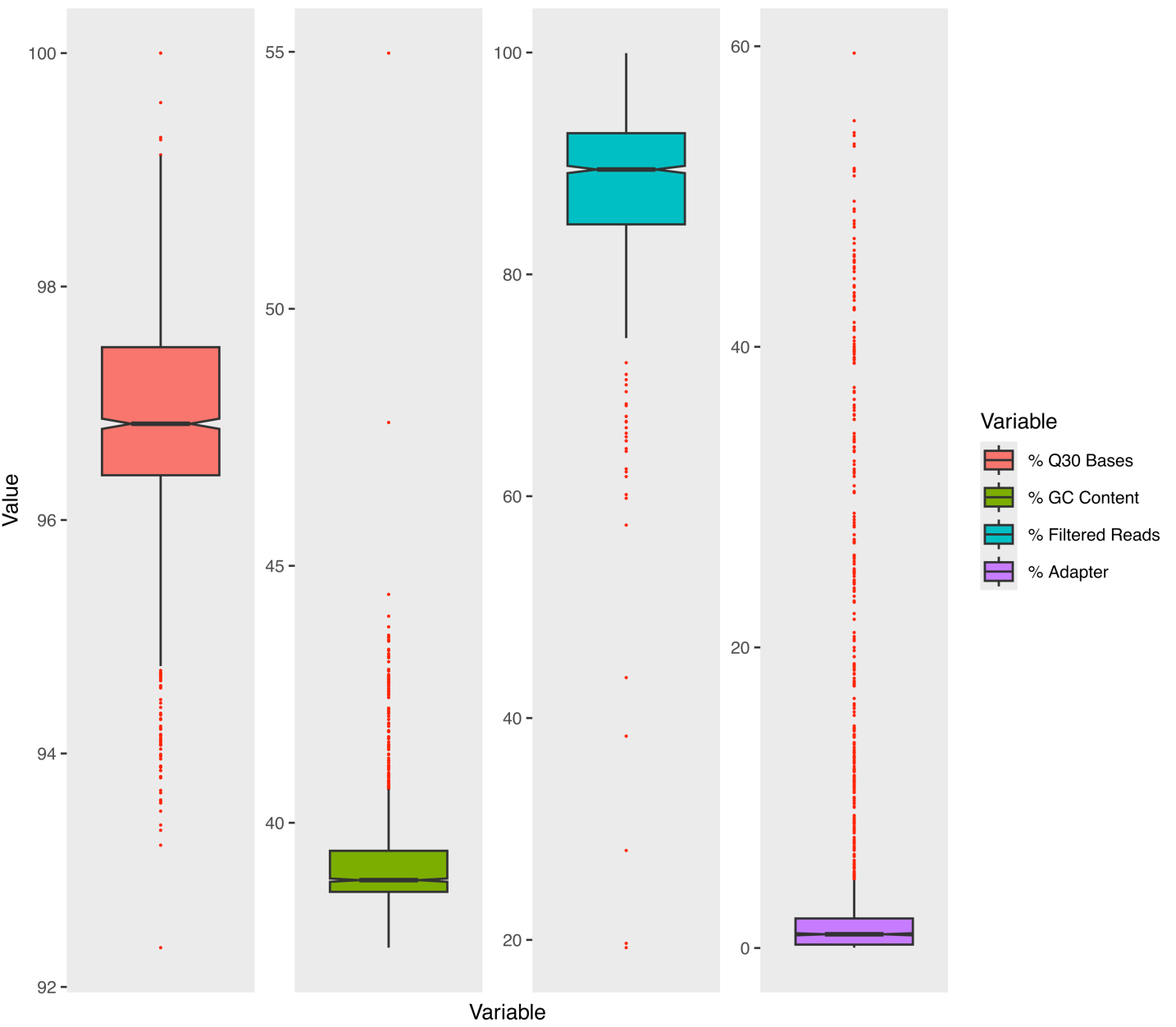
Quality metrics of raw sequence reads (n = 1,557). Q30 Bases represent the proportion of bases with Phred quality scores more than and equal to 30. Filtered reads indicate the proportion of reads retained after quality trimming. Adaptor content reflects the proportion of reads containing adapter sequence, while GC content indicates overall base composition.

**Figure S3.**
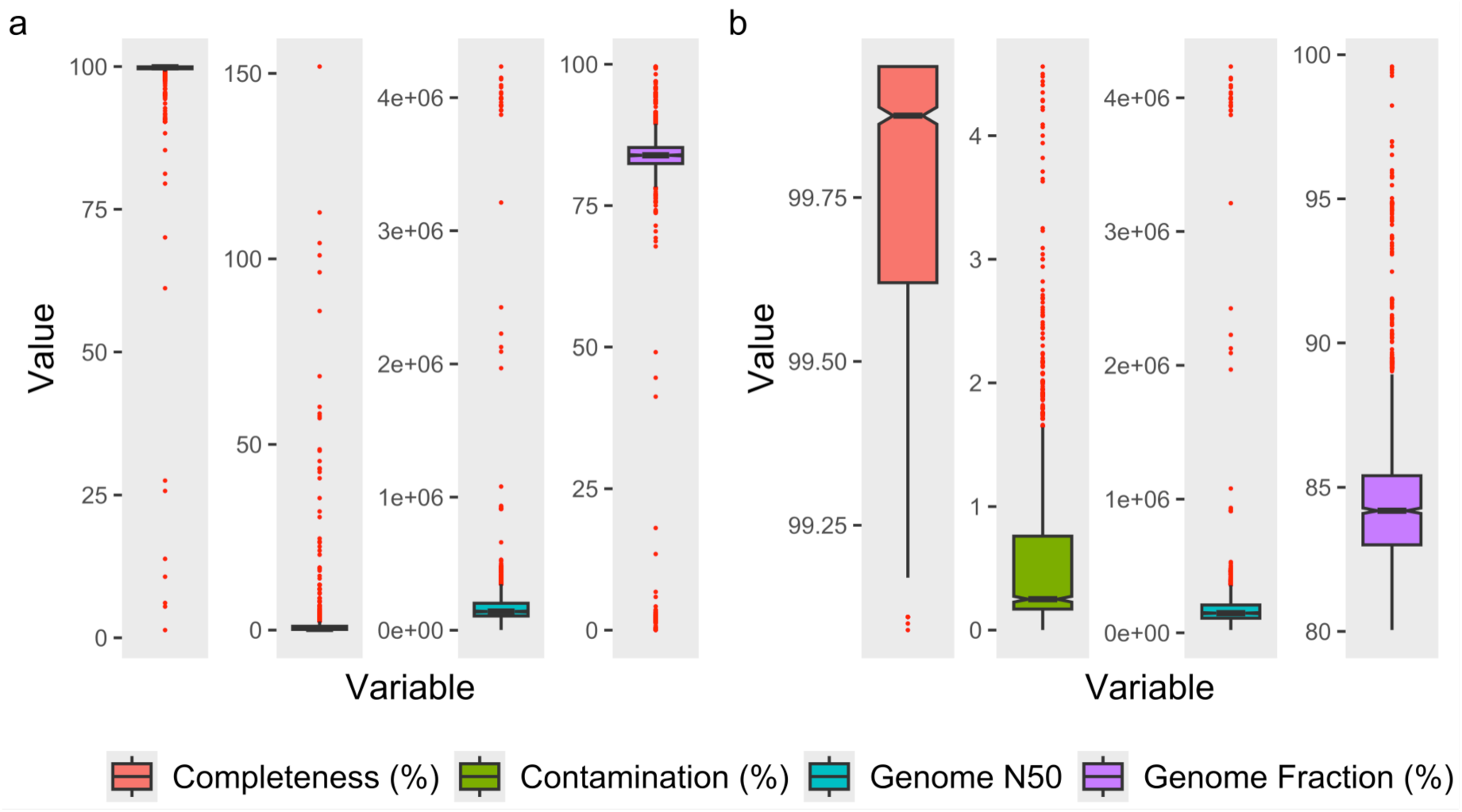
Genome assembly quality assesment. (a) Assembly quality metrics for the complete dataset (n=1,806). (b) Assembly quality metrics of the final cohort included in the GWAS (n=1,601). Assemblies were evaluated based on N50, genome coverage relative to reference, completeness, and contamination thresholds.

**Figure S4.**
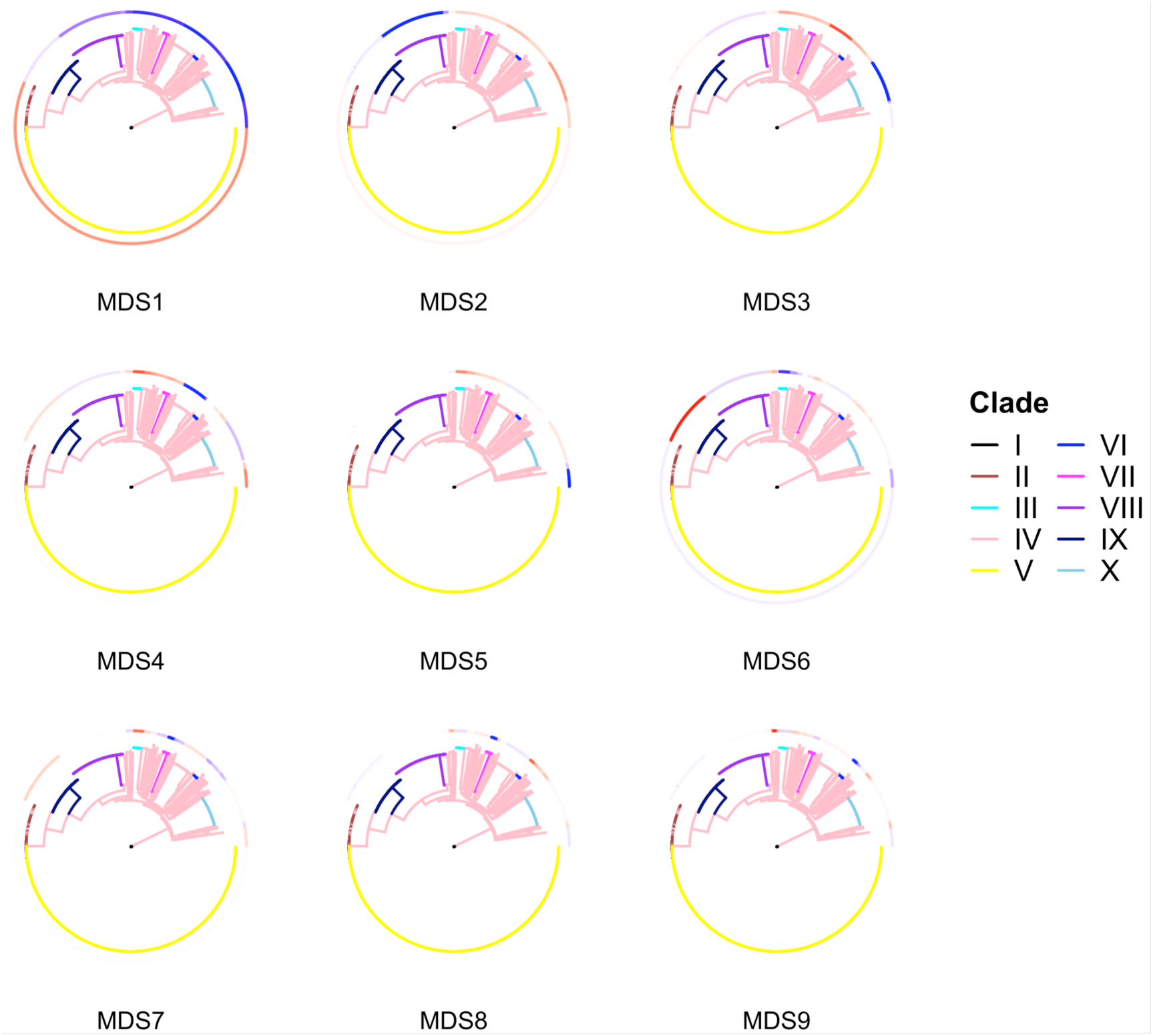
Projection of multidimensional scaling (MDS) axes onto the phylogeny. Each circular panel displays one of the first nine MDS axes, with the phylogeny colored according to BAPS-defined lineages. Clades I–X correspond to BAPS_1–BAPS_10. (BAPS, Bayesian analysis of population structure)

**Figure S5.**
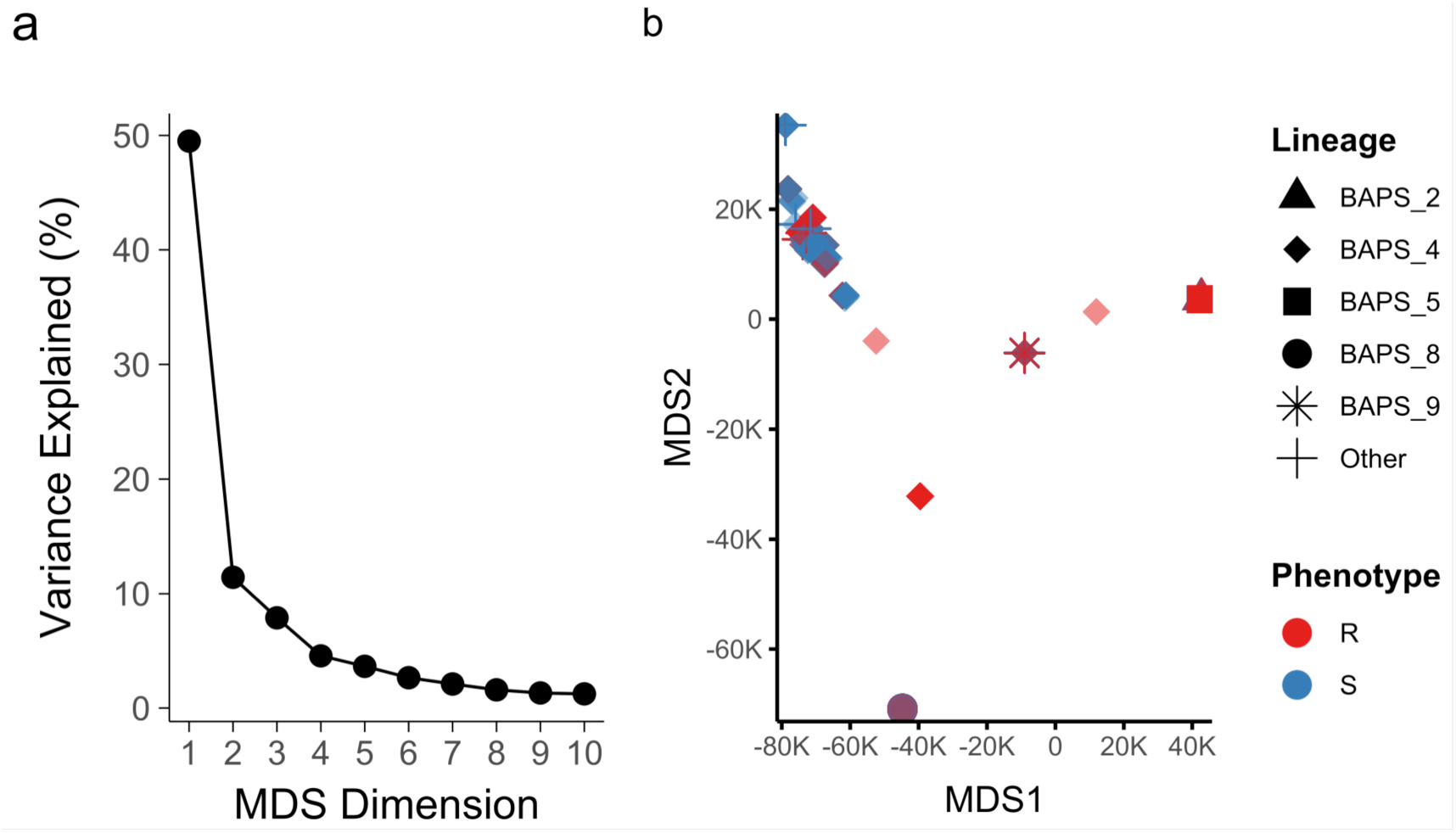
Multidimensional scaling (MDS) of core-genome SNPs. (a) Proportion of genetic variance explained by the first ten MDS axes. (b) Projection of major lineages onto the first two axes, highlighting separation of BAPS_8 and BAPS_9 from the joint BAPS_2 & BAPS_5 cluster. (BAPS, Bayesian analysis of population structure)

**Table S1.**
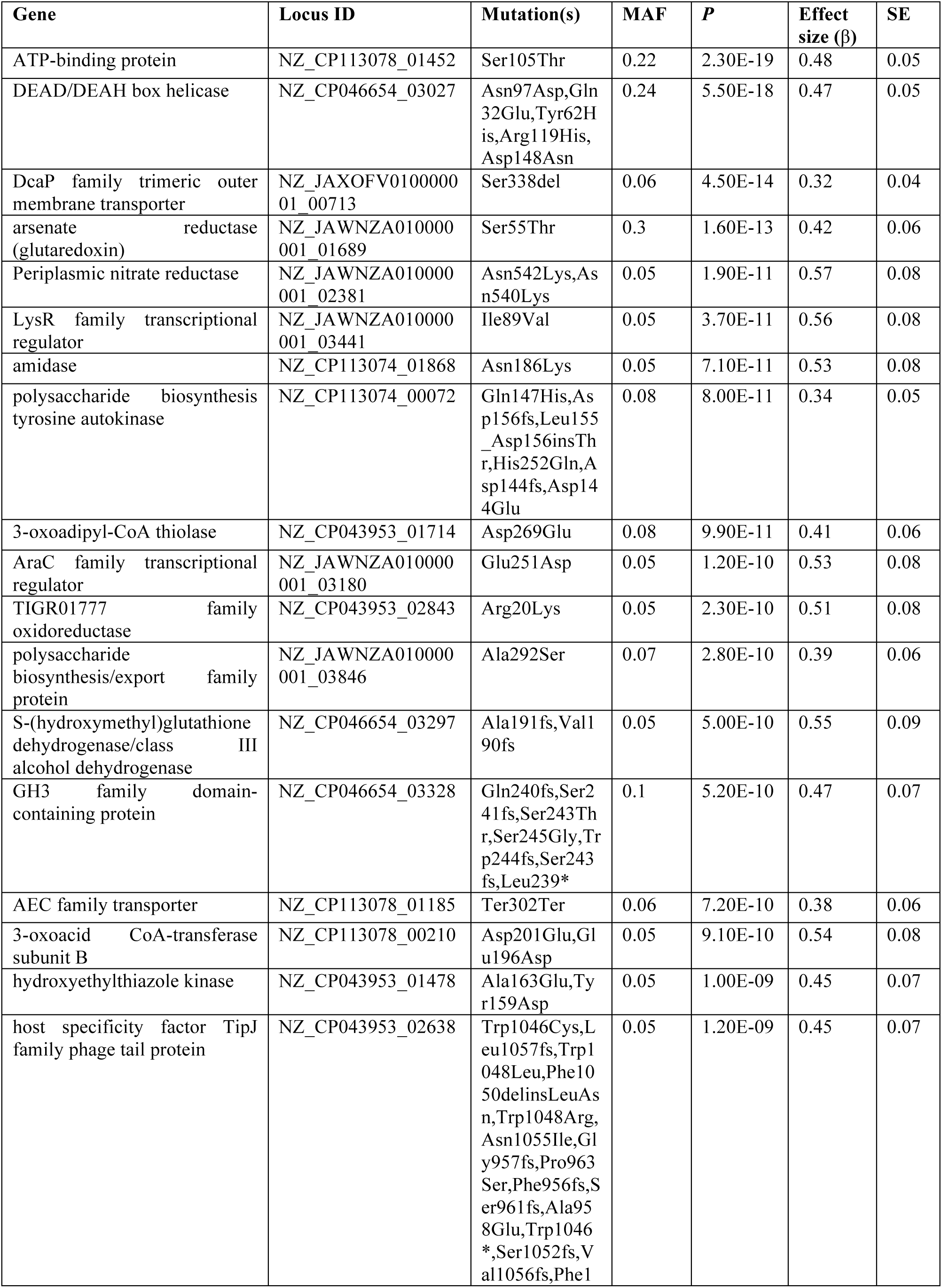
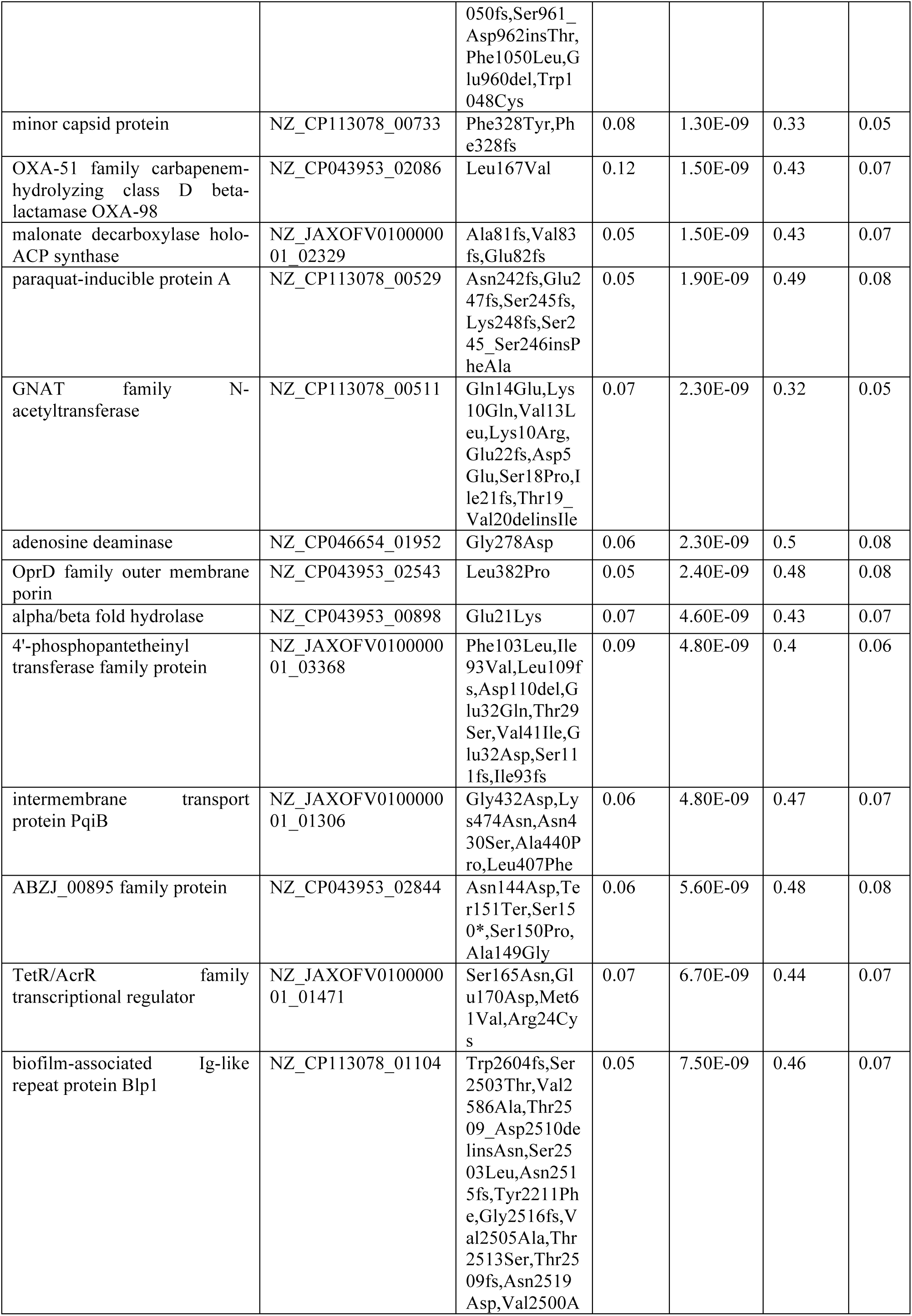
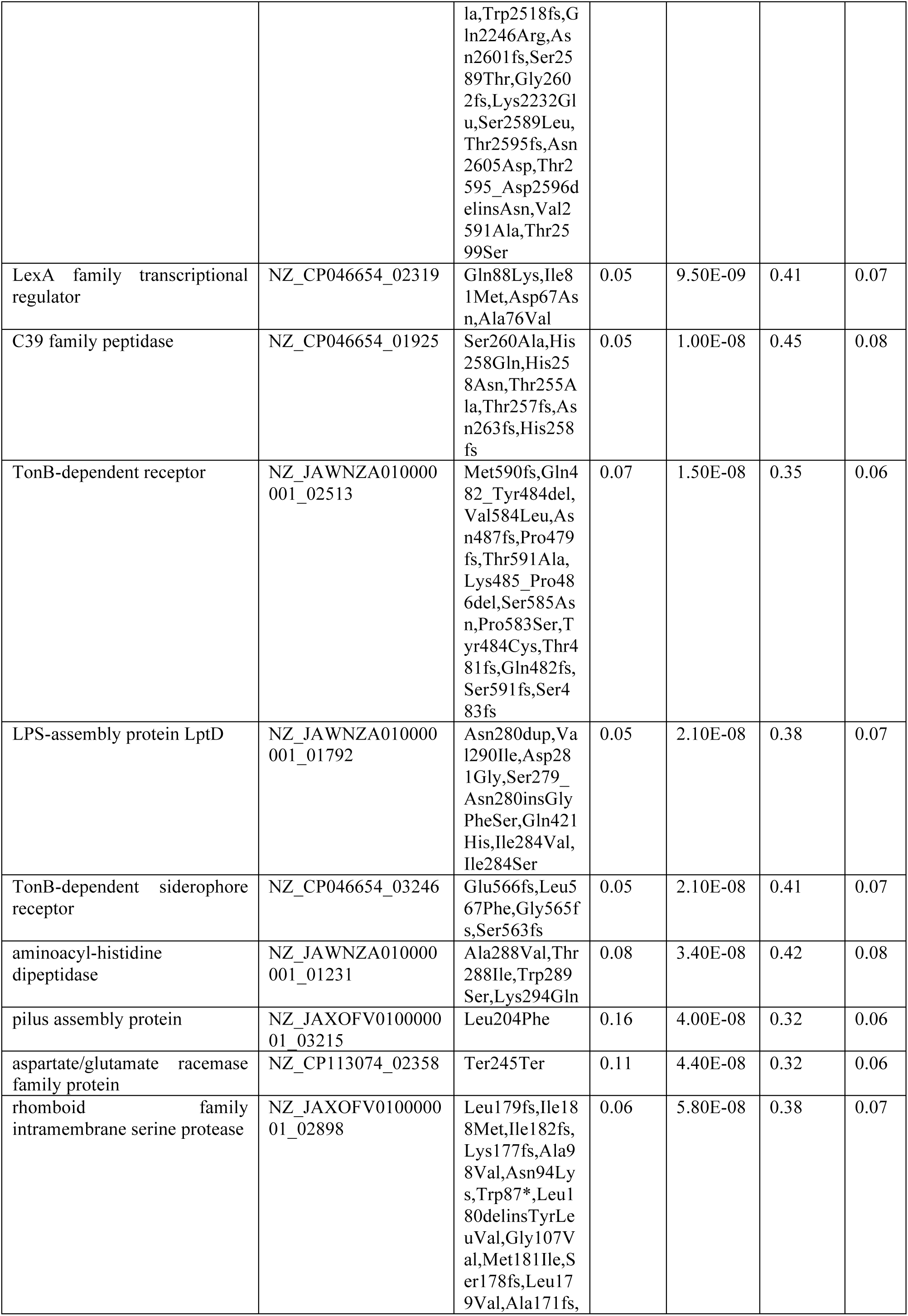
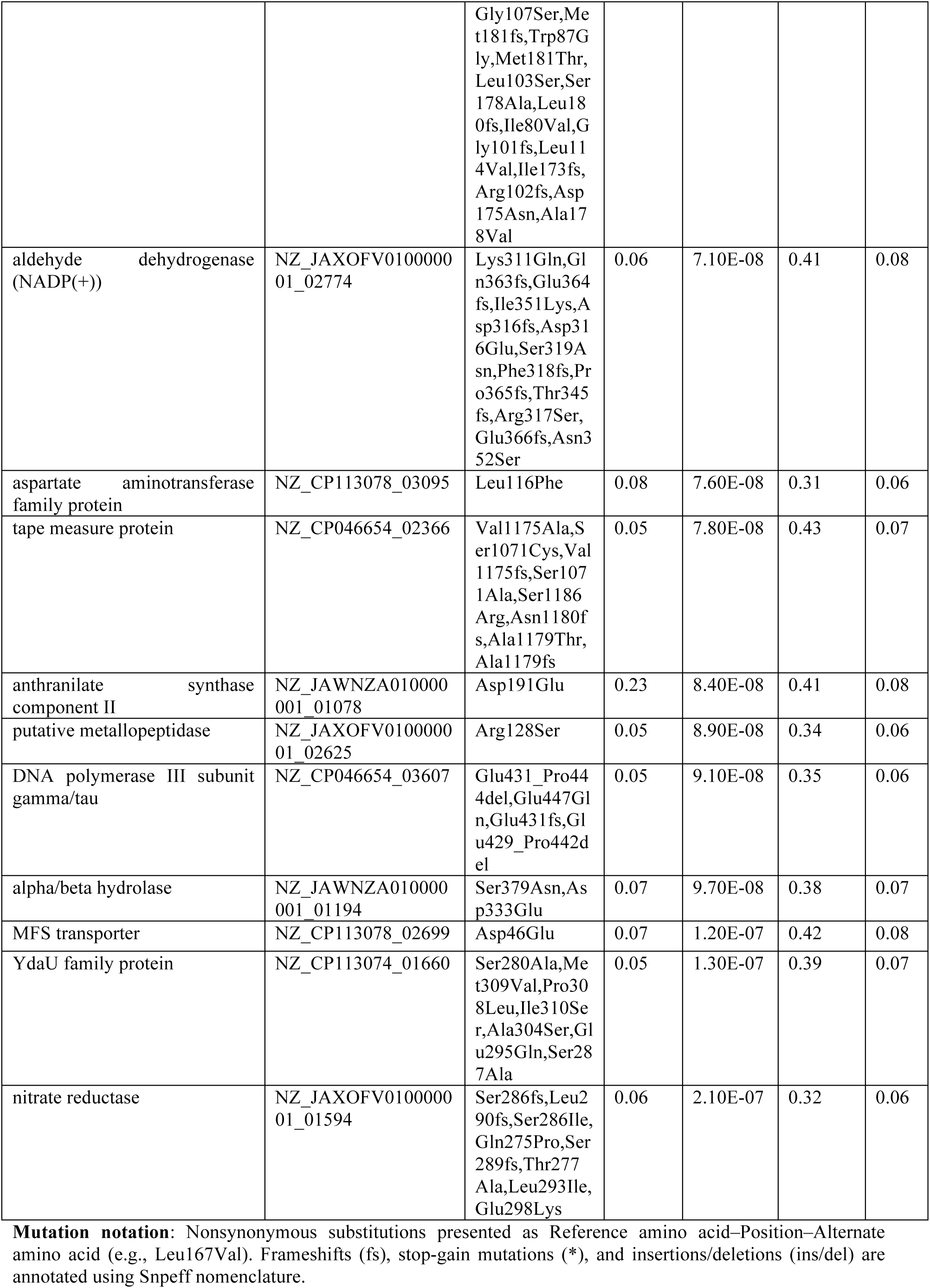
Genetic variants significantly associated with the carbapenem resistance in *Acinetobacter baumannii*, together with minor allele frequency (MAF), effect size (*β*), standard error (SE), and *P* values.

